# Deciphering the microbial contributors to methane cycling in coastal wetlands

**DOI:** 10.64898/2026.04.30.719719

**Authors:** Sophie K. Jurgensen, Djennyfer K. de Melo Ferreira, Robert Bordelon, Diana Taj, Ikaia Leleiwi, Jared B. Ellenbogen, Bridget B. McGivern, Sergio Merino, Gil Bohrer, Eric Ward, Mikayla A. Borton, Kelly C. Wrighton, Jorge A. Villa

## Abstract

Although wetlands are increasingly recognized as important contributors to the global methane budget, the microorganisms and processes involved methane cycling are poorly characterized, particularly in coastal brackish and saline systems. Here, we investigated microbial and geochemical factors contributing to methane dynamics in three coastal wetlands with different salinities, dominant vegetation types, and soil chemical characteristics. These included a freshwater flotant marsh, a cypress swamp, and a mesohaline salt marsh. Specifically, we paired methane porewater concentrations, surface fluxes, geochemistry, and 16S rRNA gene sequencing to address how microbial community composition links to porewater concentrations and its potential effects on emissions. We found that porewater methane concentrations across sites were the highest in the swamp, followed by the salt marsh and the flotant marsh, and were explained by methanogen richness and abundance. While methane-cycling microbial communities were significantly structured by salinity, two microbial taxa (*Methanosaeta* and *Methanomicrobiaceae*) were present across all sites. Hydrogenotrophs were the most abundant methanogen functional group, with *Methanomicrobiaceae* and *Methanobacterium* discriminant among wetlands. In contrast, methanotroph functional types varied among wetlands. Type I dominated the freshwater flotant marsh, while the anaerobic methanotrophic archaea the saltwater marsh. These findings contribute to an enhanced understanding of the microbiological contributions to methane emissions from coastal wetlands.

**Scientific Significance Statement Topic:** This study provides critical insights into the microbial and geochemical controls on methane emissions across coastal wetlands along a salinity gradient. Challenging the prevailing paradigm, methane porewater concentrations did not inversely correlate with salinity, as the swamp site with intermediate salinity exhibited the highest concentrations. Methanogen richness and abundance emerged as strong predictors of methane concentrations, while methanotroph richness had no predictive value. Two core methanogens, *Methanosaeta* and *Methanomicrobiaceae*, were consistently present across all wetland types. The findings highlight potential role of the water column as a biological methane filter, especially in saline environments. This study significantly advances the understanding of methane cycling in coastal wetlands by decoupling methane emissions from salinity gradients and emphasizing the role of microbial communities and local environmental factors. These insights are essential for refining biogeochemical models to forecast greenhouse gas emissions under sea-level rise and saltwater intrusion scenarios.

**Scientific Significance Statement Outlet:** This work integrates microbial ecology, geochemistry, and ecosystem structure to address interdisciplinary questions relevant to the limnological community. Here, we reveal how microbial community composition, rather than salinity alone, predicts methane emissions, offering a fresh perspective into carbon cycling in estuarine and coastal environments. The findings also provide critical insights for improving greenhouse gas emission models to predict climate change feedback from coastal areas.

## 1. Introduction

Wetlands are significant contributors to the global carbon cycle. Despite occupying just 8% of the land surface, wetlands store 30% of global soil carbon (Nahlik and Fennessy 2016) and provide ecosystem services, including wildlife habitat and flood control. This stored carbon is vulnerable to microbial decomposition, resulting in the emission of greenhouse gases including carbon dioxide and methane: in fact, wetlands are responsible for one-third of global methane emissions (Saunois et al. 2020). Coastal wetlands encompassing freshwater marshes, swamps, salt marshes, mangroves, and seagrasses along a freshwater-saltwater continuum account for 6 – 22% of the extent of wetlands globally and contribute to ∼4% of total wetland methane emissions (∼6 Tg CH_4_ yr^-1^) (Howard et al. 2017). However, there is considerable uncertainty in this estimate, given modeling constraints associated with the representation of biogeochemical processes under the dynamic redox, pH, and salinity conditions of coastal wetlands affecting methane production and consumption (O’Meara et al. 2024). The vulnerability of coastal wetlands to accelerated climate warming and sea level rise only increases uncertainty in predicting emissions from these ecosystems (Liu et al. 2019).

Microbial production and consumption drive wetland methane cycling (Bridgham et al. 2013). The temporal dynamics of these opposing processes are determined by microbes’ temperature sensitivity and hydrological dynamics to drive the seasonality in methane emissions (Chadburn et al. 2020). In brief, the majority of methane is produced by the reduction of hydrogen and carbon dioxide (hydrogenotrophic pathways), acetate (acetoclastic pathway), or methylated compounds (methylotrophic pathway) by anaerobic methanogenic archaea (Conrad 2020). Historically, the hydrogenotrophic and acetoclastic pathways are better characterized in saturated soils (Oren 1999), though methylotrophy has garnered recent attention (Vanwonterghem et al. 2016; Narrowe et al. 2019; Yuan et al. 2019; Ellenbogen et al. 2023).

To date, few studies investigating methanogenesis in saturated soils include the contributions of all three pathways (Woodcroft et al. 2018). Much of this biogenic methane diffuses up to surface soils, where it can be oxidized to carbon dioxide by anaerobic methanotrophic archaea (ANME) (Kevorkian et al. 2021) or aerobic methanotrophic bacteria (Broman et al. 2020). Biological methane oxidation in wetland soils and overlying water accounts for the oxidation of 10-90% of biogenic methane, with estimates varying considerably among wetland types and geography (White et al. 2023). Compared to wetland sediments, methanotrophs in oxygenated overlaying water columns are understudied, and thus, the methane buffering capacity of this compartment remains largely uncharacterized. In land surface models, the representation of these methane cycling processes, which ultimately determine net emissions, includes proxies for the growth and dormancy of methanogens and process-explicit reactions based on microbial functional groups (Chadburn et al. 2020). However, this approach requires information on the prevalent and dominant taxa, their distribution in the soil and water column, and environmental factors that affect them, including temperature and salinity.

Here, we investigated methane-cycling microbial communities in three wetlands along a salinity gradient featuring ecosites characteristic of the coast of Louisiana, USA, in the Northern Gulf of Mexico. We addressed how community composition (i.e., producers and consumers) links to methane porewater concentrations and its significance to methane emissions. Accordingly, we coupled greenhouse gas porewater and flux measurements, geochemical analyses, and 16S rRNA gene profiling of soil and water samples collected in October 2021. Our results suggest that ‘rules’ regarding the biogeography of methanogens and methanotrophs need to be re-examined across wetland types and ecosites. Further, our observations provide more evidence of the importance of methanotrophy as a nature-based solution to methane emissions.

## 2. Materials and Methods

### 2.1 Study site

Samples were collected from three wetland types common to the northern Gulf of Mexico: a freshwater flotant marsh (0.1-0.2 ppt, 29°85’87″ N, 90°28’64″ W), a swamp (0.6-0.8 ppt, 29°80’18″ N, 90°11’02″ W), and a mesohaline saltwater marsh (6-13 ppt, 29°49’40″ N, 89°91’52″ W) in Barataria Bay, Louisiana (Fig. 1A-B, Table S1). The freshwater (US_LA2) and saltwater (US_LA3) marshes are part of the Ameriflux network, and the swamp is within the Jean Lafitte National Historical Park and Preserve. Samples were collected between October 26 and 29, 2021 during annual peak methane emissions (Holm et al. 2016), targeting prevalent ecosites or ecohydrological patches, defined as locations with similar hydroperiod and dominant plant species. The freshwater flotant marsh contained two ecosites, one co-dominated by *Sagittaria lancifolia* and *Typha latifolia* (Sag/Typha), and the other containing solely by *Sagittaria lancifolia* (Sag). The swamp canopy was comprised of Bald Cypress (*Taxodium distichum*) and Swamp Tupelo (*Nissa biflora*), while the understory lacked clear ecosites. Thus, we sampled hummocks and hollows to account for hydrological patch heterogeneity. The saltwater marsh contained three ecosites, either dominated by *Juncus roemerianus* or *Spartina alterniflora* or open water without emergent vegetation.

**Figure 1.**
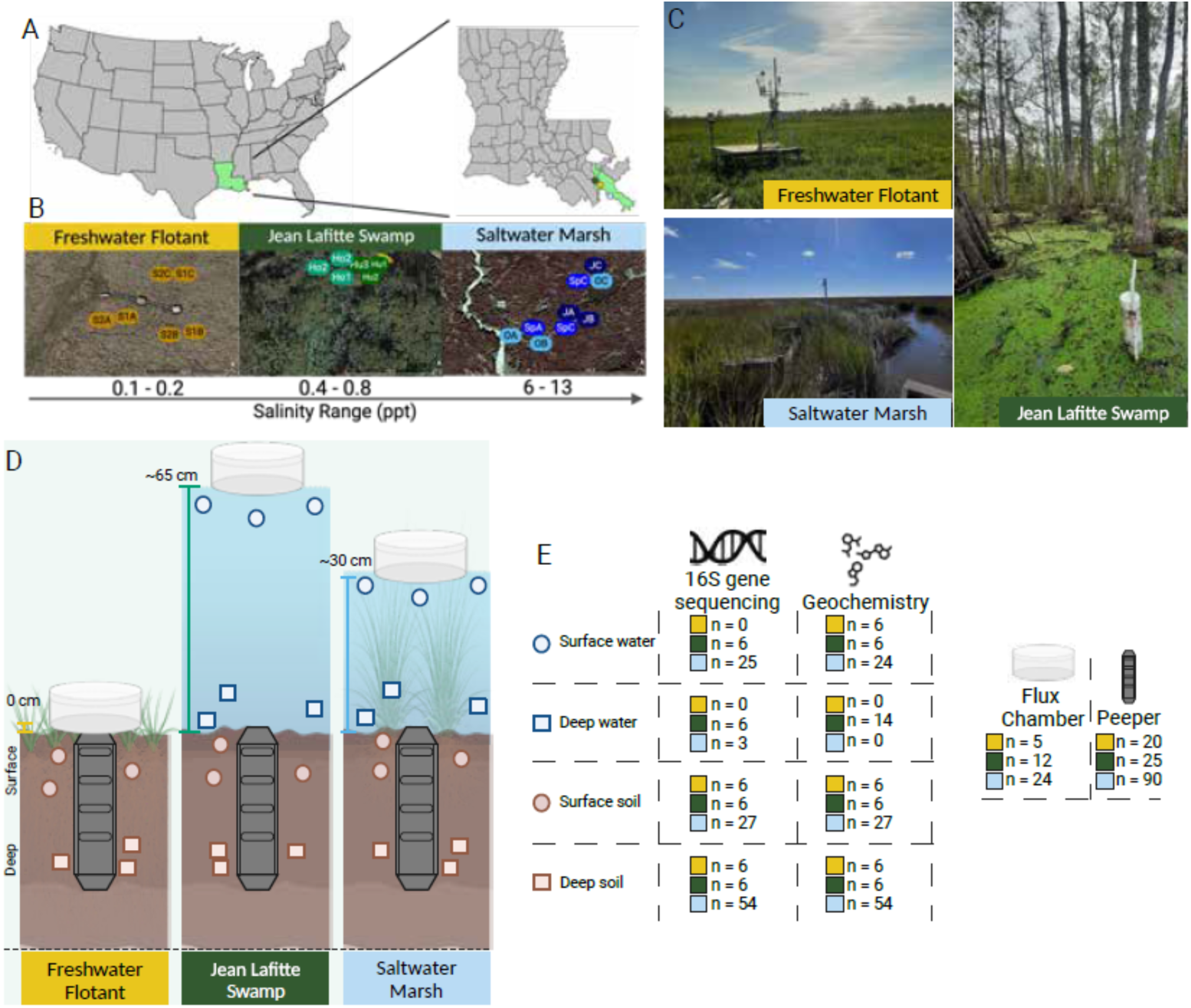
Overview of sample sites and experimental design. Map of coastal wetlands (A), ecosite locations within each wetland (B), and general site images (C) samples as a part of this study. Experimental design (D) displays sample types and collection depths, denoted by color and shape, respectively, and the average water column depth at each site. The number of microbial and geochemistry samples, as well as greenhouse gas measurements collected per wetland, compartment, and depth are displayed in (E). Created in https://BioRender.com

### 2.2 Sample collection

At each wetland, we sampled three 5×5 m plots per ecosite (21 plots in total, Fig. 1B-C). Water was collected using sterile nuclease-free 50 ml collection tubes in triplicate at the surface (∼5 cm below the water surface) and near the sediment-water interface. The swamp and salt marsh ecosites featured water tables above the soil surface, while the freshwater flotant marsh remained buoyant upon water level fluctuations, maintaining the water table below the soil surface. Hence, surface water at the freshwater ecosites was not collected, and samples at the sediment-water interface were collected with sippers that were flushed thoroughly before samples were collected and not used between sampling sites. Soil samples at each ecosite were collected using a Macaulay peat corer (5.1 cm I.D.) from which subsamples were carefully placed in sterile Whirl-Pak bags using instruments that were sterilized between sampling events using 70% ethanol. “Surface” samples were collected from the top 5 cm, and “deep” samples from 20-30 cm. At the freshwater ecosites, we were able to sample only down to 15 cm due to the high porosity of the floating mat and thus considered samples from the 10-15 cm depth range as the "deep" samples (Fig. 1D). Samples were stored in coolers with dry ice and shipped the same day of collection, then stored at −80 °C until processing. This sampling produced 145 microbial samples and 155 geochemistry samples (Fig. 1E).

### 2.3 Methane porewater concentration and flux measurements

*In situ* porewater dialysis samplers ‘peepers’ (MacDonald et al. 2013, were installed in each plot as part of another project and were sampled for this study (Table S2). We sampled every other cell (∼ every 5 cm) down to 50 cm following the protocol described in Bordeon et al. (2023). Once in the lab, methane porewater concentrations were measured from a 2 mL headspace subsample using an optical spectroscopy gas analyzer (G4301, Picarro, Santa Clara, CA) using the procedures described in Villa et al. (2024). The headspace was created from a 5 mL subsample of peeper water and 15 mL nitrogen headspace in 30 mL syringes that were agitated with a mechanical shaker for 5 minutes. The headspace (∼15 mL) was then injected into 10 ml standard chromatography sealed glass vials that had been previously evacuated. These headspace samples were swiftly analyzed for methane concentration within the following 24 hours.

*In situ* methane fluxes were measured with the same gas analyzer used for porewater gas analyses (G4301, Picarro, Santa Clara, CA). Three chambers were set within 0.5 m of peepers at each sample plot (Fig. 1E). At the marsh sites, collars (30 x 30 x 15 cm) were pre-installed to allow time for equilibration. Then, transparent chambers (100 or 120 cm tall) equipped with 12V fans to ensure gas circulation were attached to these collars for 3-minute deployments. At the swamp, where the water level exceeded the collars’ height, we deployed cylindrical transparent chambers (0.066 m^2^ area, 0.012 m^3^ volume) with attached foam for floatation. The time series of methane concentrations from each chamber deployment was transformed into flux using chamber dimensions and field temperature and pressure, following the fit of a non-linear approximation as described in Villa et al. (2021). Reported fluxes correspond to methane emissions from the wetland surface comprising the soil and water column, including vegetation where present. We used only fluxes where the diffusive and ebullitive signals were clearly discernible and had r2 > 0.7 fits for the diffusive phase. Of 41 attempted flux measurements, 23 passed this quality control (Table S3).

### 2.4 Geochemical measurements

Salinity concentrations were obtained from the Coastwide Reference Monitoring System (CRMS, https://lacoast.gov/crms/#). We used 2021 data from stations 3166, 0234, and 0224 stations representing the freshwater flotant marsh, swamp, and saltwater marsh, respectively. Dissolved oxygen and temperature were measured *in situ* using a fiber optic oxygen meter (PreSens Fibox 4) at the saltwater marsh site (Fig. S1). Redox potential (ORP) was also measured *in situ* using a portable Mv meter and redox electrode (Faulkner et al., 1989) (Fig. S2). Ion chromatography (Dionex™ ICS-6000 DP) was used to measure anion (fluoride, acetate, chloride, nitrite, bromide, nitrate, sulfate and phosphate) and cation (ammonium, calcium, magnesium, potassium, sodium and lithium) concentrations, while Fe (II) was measured using Hach FerroVer™ Iron Reagent Packets (Fig. S3). Nitrite, lithium, and phosphate were not detected in any samples and were not included in the following analyses. Soil and water pH were determined using the Fisherbrand™ accumet™ AB150 pH benchtop meter (Fig. S4, Table S4).

### 2.5 DNA extraction, 16S rRNA gene sequencing and processing

Total nucleic acids were extracted from soil and water samples using the Zymo Research Quick-DNA™ Fecal/Soil Microbe Microprep Kit, following the manufacturer’s and other standard extraction protocols, including blanks in each round of extractions. The blanks were not amplified/sequenced because they were all below the limit of detection (i.e., 20,000 reads) and did not amplify via PCR. DNA concentration was determined using a Qubit fluorometer, and extracted DNA was stored at −80°C until library preparation. The V4 region of the 16S rRNA gene was amplified using the 515F (GTGCCAGCMGCCGCGGTAA)/806R (GGACTACHVGGGTWTCTAAT) primer set (XXXXX) following the Earth Microbiome Project PCR protocol. Libraries containing a unique sequence tag to barcode each sample were sequenced on an Illumina MiSeq using 251 bp paired-end sequencing chemistry at the Microbial Community Sequencing Lab (University of Colorado Boulder). Total reads were demultiplexed and analyzed using QIIME2 (2021.2.0, Bolyen et al. 2019) with the DADA2 pipeline (Callahan et al., 2016) and clustered at 99% identity to yield amplicon sequence variants (ASVs) that were then taxonomically classified using SILVA (silva132.250, Quast et al. 2013). As rarefaction did not affect results, samples were not rarefied; instead, samples with fewer than 5,000 reads were not retained for further analyses (Table S5). Chloroplast and mitochondria hits were filtered from the produced ASV table using QIIME2.

### 2.6 Taxonomic assignment of methanogens and methanotrophs

ASVs were manually classified as methanogens and methanotrophs via the known physiology of the finest taxonomic resolution for each ASV (Lyu and Liu 2018; Evans et al. 2019; Guerrero-Cruz et al. 2021) at the taxonomic resolution of family level and below (Table S6). Where methanogenesis and methanotrophy were not conserved across order or higher we did not consider ASVs as methane-cyclers. For example, Methylomirabilota and Methanosarcina classes (SILVA 132) contain some non-methanotrophic and non-methanogenic lineages (e.g., MBNT15 and ANME-3 genera, respectively) (Ivanova et al. 2022; Knittel et al. 2005). Members of the *Syntrophoarchaeaceae* were classified as likely alkanotrophs (Evans et al. 2019), and *Bathyarchaeia* sp. "uncultured_methanogenic" were not classified as methanogens here based on the undetermined role of encoded mcr genes (Vanwonterghem et al. 2016).

### 2.7 Statistical analysis and data visualization

Statistical analyses were performed using R (v4.2.0, R Core Team 2023) unless otherwise stated. Comparisons of methane porewater concentrations between wetlands (i.e., pooled depths and ecosites) and soil profile depths of each ecosite were determined by Shapiro-Wilks and pairwise Wilcoxon tests. Visualizations were generated with the *ggplot2* package (Wickham, 2016) and further modified using BioRender. Alpha and beta diversity metrics were determined using the *vegan* package (Oksanen et al. 2024). Correlations between environmental variables were performed using the ‘rcorr’ function of the *Hmisc* package (Harrell and Dupont, 2024) using Spearman coefficients and visualized using the *corrplot* package (Wei and Simko 2024) with an α of *p* < 0.05.

We performed a principal component analysis (PCA) of geochemical measurements (pH, salinity, anions and cations) using the ‘envfit’ function of the *vegan* package (Oksanen et al. 2024), and compared this to a non-metric multidimensional scaling (NMDS) analysis based on Bray-Curtis dissimilarity of total and methane-cycling microbial communities. Geochemical compounds driving spread in ordination were based on the Euclidean distance from the center. We used an analysis of similarities (ANOSIM) and multi-response permutation procedure (MRPP), as well as permutational multivariate analysis of variance (PERMANOVA) to compare wetland type, dominant vegetation, depth, and sample type.

Indicator species analysis was performed using the *indicspecies* package (Cáceres and Legendre, 2009) to determine genera significantly associated with each site at the total community and methane-cycling community levels. Genera with IndVal > 0.7 and p < 0.01 were considered significant. Generalized additive models (GAMs) using log-transformed microbiome data to predict log-transformed porewater methane and fluxes were created using the mgcv package (Wood, 2011). Models were fit separately for each predictor-response combination using thin plate regression splines with a maximum basis dimension of k = 5 and smoothing parameters estimated by restricted maximum likelihood, with smooth term complexity was quantified using effective degrees of freedom (edf). Heteroscedasticity of these GAMs was evaluated via Breusch-Pagan test using the lmtest package (Zeileis and Hothorn 2002).

## 3. Results

### 3.1 Methane porewater concentrations and fluxes across wetland types

Methane porewater concentrations were highest in the swamp (median; mean ± std. error) (500.8 μM; 500.6 ± 81 μM, n = 20), followed by the salt marsh (262.5 μM; 311.9 ± 28 μM, n = 90), and lowest in the freshwater marsh (27 μM; 44.4 ± 14 μM, n = 25) (p < 0.05 for all pairwise comparisons) (Fig. 2A, Table S2, Bordelon et al. 2023). In turn, methane fluxes at the flotant marsh (median (IQR)) (42.7 (26.5-58.8) µmol m^-2^ s^-1^, n = 2) were higher and had considerable larger variation than fluxes at the swamp (5.78 (2.31-16.6) µmol m^-2^ s^-1^, n = 5) and salt marsh (0.013 (0.007-0.28) µmol m^-2^ s^-1^, n = 16) (pairwise Wilcoxon p < 0.05) (Table S3).

**Figure 2.**
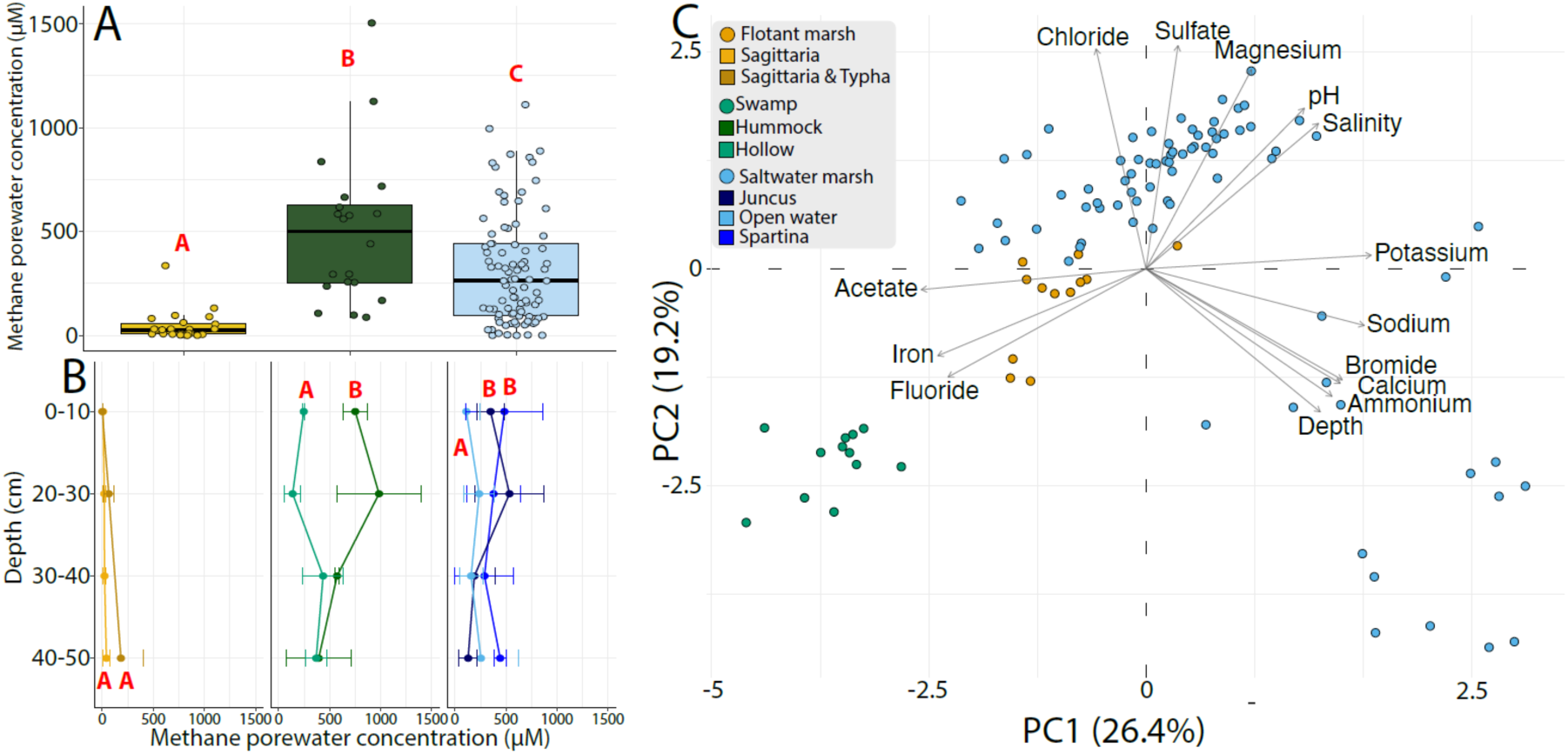
Porewater concentrations across wetlands (A), with boxplots representing the median (horizontal black line inside the box), lower and upper quartiles (25% and 75% of interquartile ranges), and outliers for each subsite. Mean porewater concentrations and standard deviations in the soil profile at the ecosite level at each wetland (B). Letters in (A) and (B) denote significant differences (pairwise Wilcoxon p < 0.05).

Ecosite-level differences in porewater concentration were observed in the swamp, with methane concentrations in hummocks exceeding those of hollows (pairwise Wilcoxon p = 0.0017, Fig. 2B). Differences were also observed at the saltmarsh, with larger methane concentrations at the *Spartina* and *Juncus* vegetated ecosites than in the open water (pairwise Wilcoxon p = 0.0014 and 0.025, respectively) (Fig. 2B). Interestingly, methane porewater concentrations in vegetated patches at the saltmarsh (*Spartina* and *Juncus*) and the flotant marsh (*Sagittaria* and *Sagittaria/Typha* mix) were not statistically different and only showed differences with depth for the *Juncus* species. The methane profile in the soil at this ecosite peaked at 20-30 cm and showed lowered concentrations at 30-40 cm and 40-50 cm (pairwise Wilcoxon p < 0.05). A principal component analysis (PCA) revealed the swamp to be most distinct from the other sites, largely due to differences in iron and fluoride concentrations (Fig. 2C).

### 3.2 Methane-cycling microbial community structure

The 105 soil and 40 water samples yielded 8,284,621 high-quality reads. We identified a total of 60,605 ASVs clustered at 99% identity, which were then taxonomically assigned using SILVA (Fig. S5, Table S5). Of these, 709 could be taxonomically identified as being methanogenic or methanotrophic. Shannon diversity (pairwise Wilcoxon p < 0.05), but not species richness (pairwise Wilcoxon p > 0.05), varied among wetlands (Fig. 3A, B). When considering only the methane-cycling communities, species richness, Shannon diversity varied significantly among wetlands (pairwise Wilcoxon p < 0.05, Fig. 3C, D), with both metrics decreasing with increasing salinity. Similarly, the relative abundance of methanogens and methanotrophs was negatively correlated with salinity, though only methanotroph abundance was significant (p < 0.05, Fig. 3E). This trend was driven by the relationship between hydrogenotrophic methanogens and salinity. Notably, ANME were only detected in the saltwater marsh, and their abundance was thus positively correlated with salinity. Furthermore, variance in soil microbial community structure both at the whole community level (Fig. 3F-G) and methane cycling community level (Fig. 3H-I) was associated with wetland, ecosite, and depth (PERMANOVA p < 0.01). Salinity and related variables (e.g., chloride (Cl^-^), magnesium (Mg), and fluoride (F^-^)) and pH were the main contributors to variation along Axis 1 in both cases suggesting increasing marine influence, while depth was the single largest contributor to Axis 2 indicating the influence of redox and mineral interactions that manifest at the ecosite level (Fig. 3F,H, Fig. S6).

**Figure 3.**
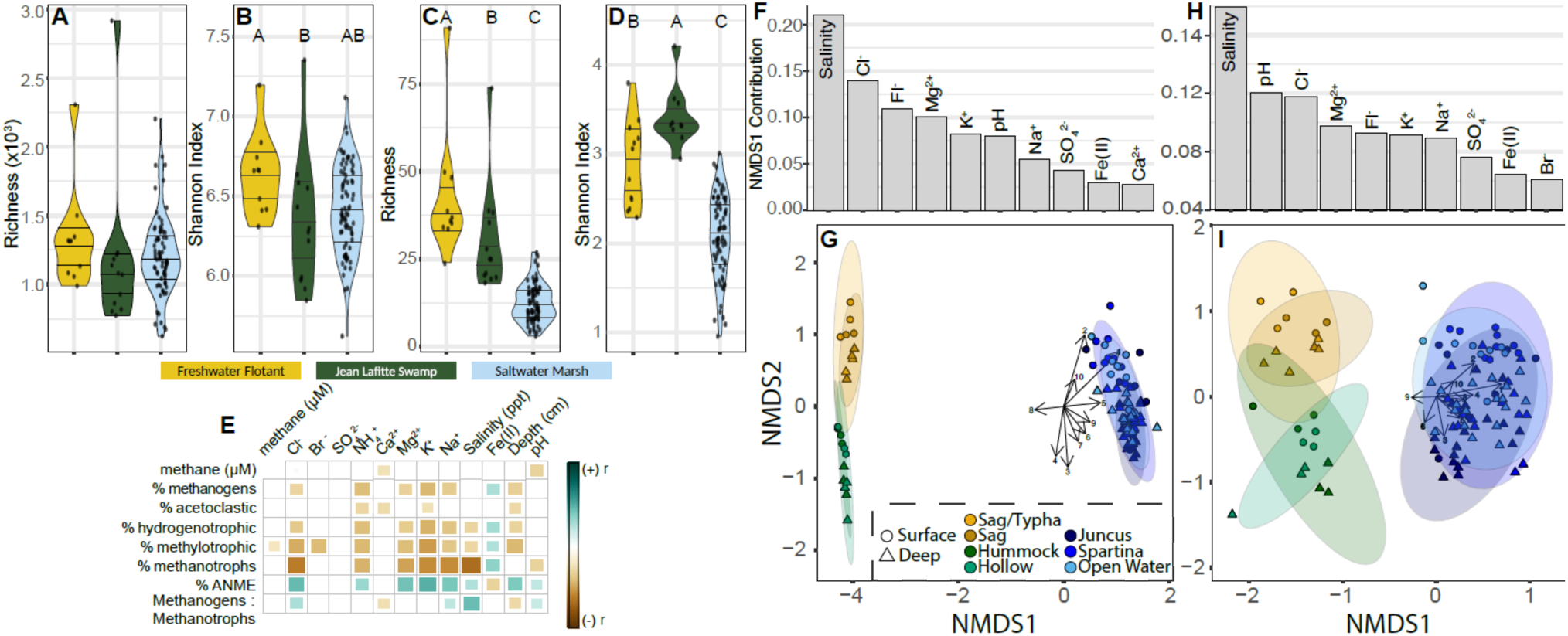
Diversity metrics and correlations with environmental variables. Alpha diversity of total microbial communities (A, B) and methane cycling microbial communities (C, D) in soils of each wetland. Horizontal lines within violin plots denvote quartiles. Points denoting each sample are jittered to improve visualization. Letters above plots denote significant differences at the wetland level (Tukey’s post-hoc test p < 0.05). Correlation matrix of environmental variables and the relative abundance of methanogens, methanotrophs, and methanogen functional groups (E). Only significant (p < 0.05) relationships are shown as boxes, with color and size denoting the degree of the correlation. Non-metric multidimensional scaling (NMDS) and associated drivers of total microbial communities (F, G) and methane cycling communities (H,I) in soils of each wetland as determined by Bray-Curtis dissimilarity. The top 10 biogeochemical compounds driving the ordinations are displayed for each (largest envt vectors). In both ordinations, NMDS2 is most strongly driven by depth. Wetland, ecosite, and depth are also significant drivers of both ordinations (PERMANOVA p < 0.01 in all cases). Salinity was measured at the wetland type level (Table S4).

Based on the differences observed in the NMDS, we next investigated the incidence of methanogens and methanotrophs among water (Fig. 4 A-B) and soil (Fig. 4 C-D) samples across sites (Fig. S7 and S8). *Methanosaeta*, an obligate acetoclast, and Methanomicrobiaceae, a hydrogenotroph (best assignment at the family level), were the only core genera (present in > 80% of all soil samples) among both surface and deep soils from each site (Fig. 4A,C). Hydrogenotrophs were the most abundant methanogen functional type across sites (Fig. 4A,C, Fig. S9), and their abundance was positively correlated with Fe(II) concentrations (Fig. 3E). Hydrogenotroph abundance was also weakly negatively correlated with sulfate concentrations (Fig. 3E, Table S7), signaling that they may be outcompeted by sulfate-reducing bacteria enriched by higher sulfate concentrations. We note this response could be because we averaged the relative abundances of many rare taxa (n=130 ASVs present in only one sample), as these rare members may drive observed patterns. All five methanogenic genera discriminant among wetlands were significantly associated with the freshwater flotant marsh and/or the swamp, but not the saltwater marsh (Table S8), reflecting their overall decrease in abundance and diversity with salinity (Fig. 3E, 4A,C).

**Figure 4.**
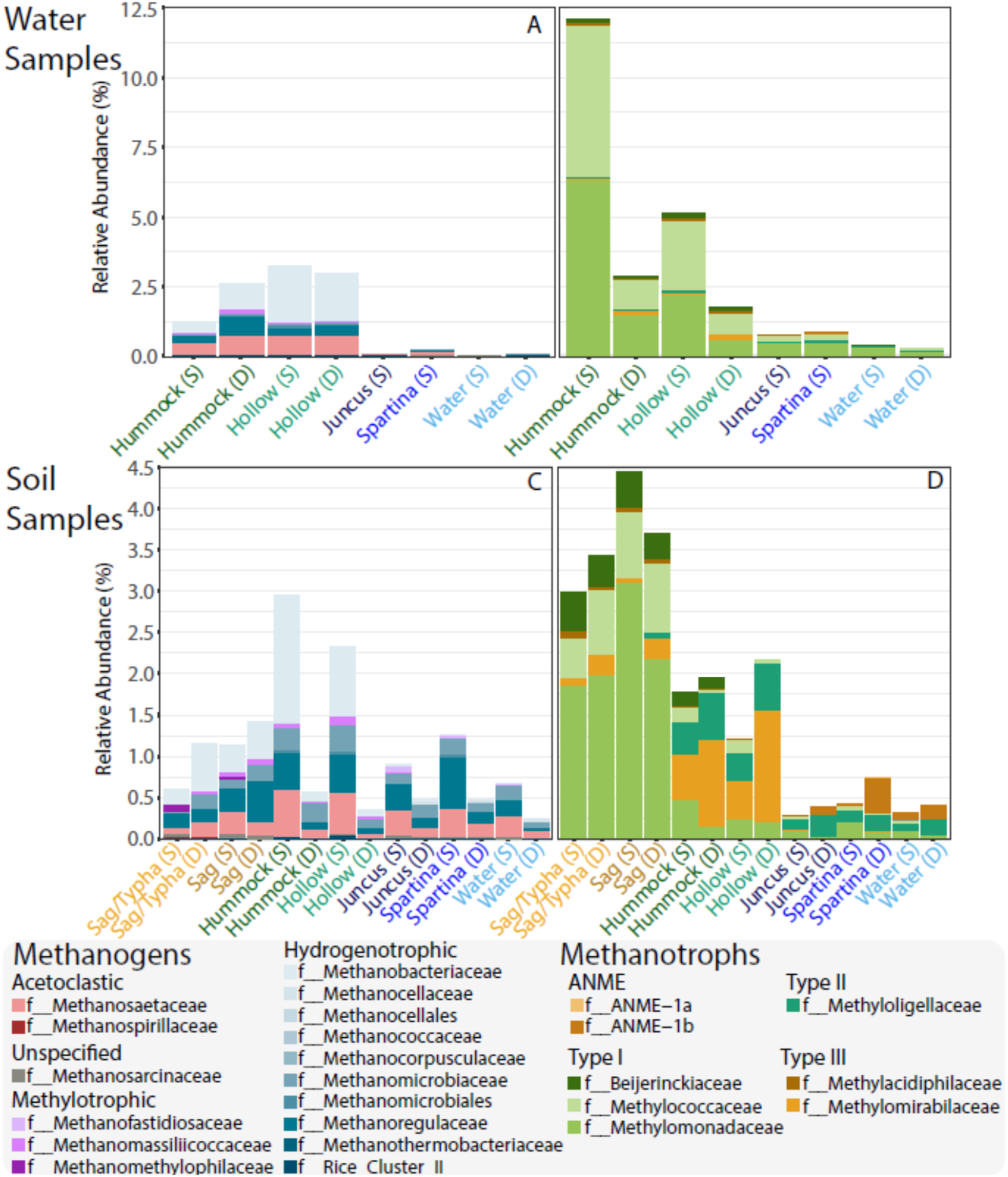
Relative abundance of methanogen and methanotroph taxa across sites, ecosites, and depth in water (A and B) and soil (C and D). Color shades denote functional guilds (acetoclasts, hydrogenotrophs, methylotrophs, unassigned substrate methanogens, anaerobic methanotrophs, Type I, Type II, and Type III methanotrophs).

The relative abundance of methanotroph communities in water and soil samples decreased from the freshwater flotant to the saltwater marsh, with water samples consistently having higher abundances of aerobic methanotrophs compared to soils (Fig. 4B,D). In addition to variations in the relative abundance of methanotrophs across wetlands, we noted that methane oxidizers exhibited more site-specific diversity compared to methanogens. Few methanotroph taxa were shared among different wetlands, and no core members were found between them (Fig. S8). Cumulatively, aerobic Type I methanotrophs, including *Methylobacter* and *Methylomonas* were dominant (Fig. 4B,D), accounting for up to 90% of the relatvive abundance of methanotrophs in some samples. However, Type II methanotrophs including Methylogellaceae and *Methyloceanibacter* were markedly more abundant in the swamp and saltwater marsh. Notably, methanotrophs of the Verrucomicrobiota were almost exclusively present in swamp soils and anaerobic methanotrophs (ANME) were only present in the saltwater marsh, representing dominant and discriminant members of those soil methanotroph communities (Fig. 4D, Table S6). Though their abundance and diversity decreased along our salinity gradient, diverse methanotroph types were detected across sampled wetlands.

### 3.3 Predicting methane concentrations and fluxes

Across sites, we observed i) porewater methane concentration differences (Fig. 2A), ii) distinct abundance patterns of methane cycling microbes (Fig. 4, Fig. S6), and iii) weak correlations between porewater methane concentrations and the relative abundance of methane cycling community members (Fig. 3E). To determine whether methane cycling taxa were appropriate predictors of in situ methane concentrations and methane fluxes, we utilized general additive models (GAMs). Fluxes were strongly predicted by methanogen and methanotroph richness and abundance but not the ratio of methanogens:methanotrophs (approximate Wald p < 0.05 for all). Notably, porewater methane concentrations were strongly negatively related to methanotroph abundance and richness, but not methanogens (Fig. 5). In all cases, the relationship between methanotroph predictors and methane responses was not linear (edf > 1), while methanogen-based predictions were. While stronger relationships were observed between these predictors and methane fluxes for water samples alone (Fig. S10, Table S9), these may be spurious due to confounding from higher methane production in the soils.

**Figure 5.**
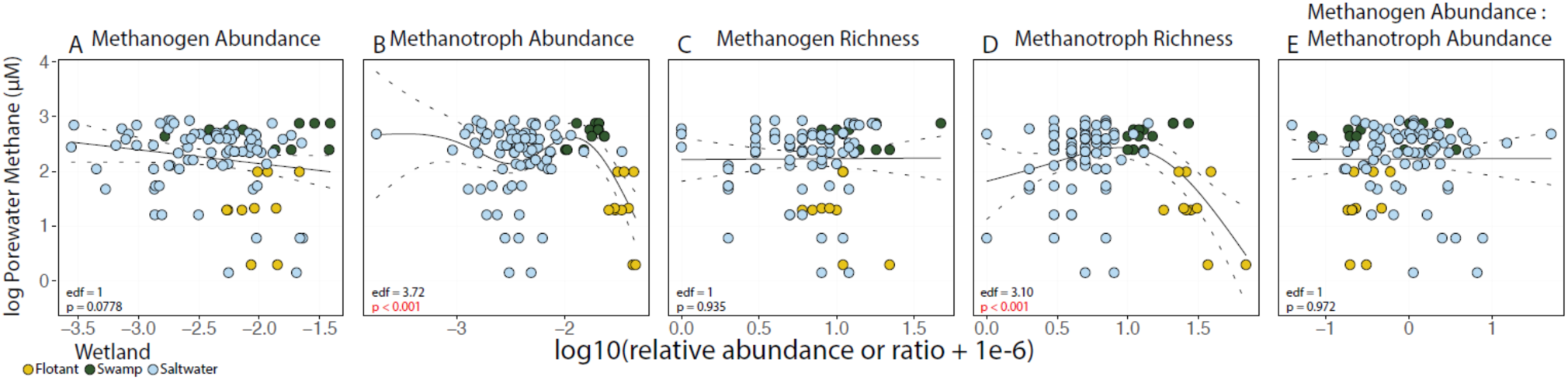
Generalized additive model (GAM) fits of methane porewater concentrations as a function of methanogen (A) and methanotroph (B) abundance, methanogen and methanotroph richness (C,D), and ratio of methanogens: methanotrophs abundances (E).

## 4. Discussion

### 4.1 Methane porewater concentrations do not follow a salinity gradient

The observed differences in methane concentrations across sites and varying geochemistry associated with them support the idea that the freshwater-saltwater continuum is multidimensional and methane dynamics can be decoupled from salinity as the single controlling variable by the combined effects of sulfate and other ions, redox conditions, and vegetation. For instance, sulfate concentrations may vary independently of salinity due to localized sulfate depletion, leading to elevated porewater methane concentrations even when salinity is high (Poffenbarger et al. 2011). Besides, salinity can be driven by other ions or other inorganic chemical compounds different from sulfate, like nitrate or ammonium, or elements such as chloride, sodium, and potassium (Noe et al. 2013), which are also present at our swamp site (Jurgensen et al., 2025).

In addition, methane porewater concentrations in the flotant marsh were low compared to other freshwater coastal marshes. Studies in coastal wetlands in the US Atlantic coast and Southeast Asia report values of methane concentrations that are larger than 100 μM at low salinities (i.e., ∼ 0.1 ppt) (Poffenbarger et al. 2011), or almost twice our mean. Low porewater concentrations of greenhouse gases in coastal wetland substrates, such as the floating vegetated mat we sampled at this site, have been explained by high rates of lateral flows of dissolved methane and ebullition (Tong et al. 2010). These two processes are enhanced by the high porosity, which is characteristic of the flotant marsh substrate and its high organic matter content (Wright et al. 2018). The freshwater marsh is further hydrologically connected to Barataria Bay and thus subject to storm surges during major storms that introduce saltwater and, with it, sulfate, explaining its relatively high values at the freshwater ecosites (Table S4), and to some extent, a potential inhibitory effect on methane production.

Beyond the controlling effect expected from salinity over larger scales, methane concentration at the ecosite level is the result of the balance between inputs, including *in-situ* production, and lateral and vertical fluxes in the dissolved phase, and outputs, including methane consumption, lateral flux, and emissions to the atmosphere through diffusion, ebullition, or plant-mediated transport. These processes are, in turn, modulated by ecosite-specific environmental conditions. Amongst the more studied are the water levels (and resulting redox conditions), temperature, availability of electron acceptors and substrates for organic matter decomposition, plant phenology, soil physicochemical characteristics, and the structure of the microbial community in the soils (Treat et al. 2007; Segarra et al. 2013). For example, we found larger methane concentrations in the hummocks than in hollows (Fig 2B), contrary to the current paradigm based on studies from northern peatlands (Krohn et al. 2017), but echoing a recent study at another cypress swamp, which attributed the increased concentrations to the predominance of cypress tree root structures (i.e., knees) with enhanced methane transport, which are more abundant in hummock spots (Ward et al., 2020). Further, differences between vegetated and open water ecosites observed in the saltwater marsh (Fig. 2B) may be explained by emergent vegetation, which modulates methanogenesis and fluxes to the atmosphere. Vegetation enhances methane production by providing organic matter through leaf litter turnover and exudates, which serve as carbon sources for microorganisms during decomposition and methanogenesis (Hatala et al. 2012). However, vegetation also transports oxygen to the rhizosphere, limiting methanogenesis (Haviland and Noyce 2024) and can also act as a conduit for methane, thus reducing the resistance to methane transport and, thus, reducing the concentration gradient from the soil to air and ultimately lowering the pool in the soil (Bansal et al. 2020). Our time-limited but heterogeneous snapshot highlights the complexity of methane dynamics in coastal wetlands, showing that while salinity can have an overarching influence on methane production, local site-specific factors, modulated by vegetation, soil or sediment structure, and microtopography, may ultimately shape methane concentrations and fluxes.

### 4.2 Methane-cycling microbial communities are structured by wetland type and depth

Inhibitory effects of salinity are reported to control the distribution of methanogens and to suppress methane production across estuarine wetlands, even at lower salinity levels like those presented here (Webster et al. 2015; Luo et al. 2019). This is typically associated with the direct toxicity of the salts or with an abiotic effect of salinity, which solubilizes ionically bound ammonium and is a known inhibitor of methanotrophs in wetlands and lake sediments (Hartman et al. 2024). In our study, salinity, chloride and ammonium were significantly negatively correlated with methanogen and aerobic methanotroph abundance, though ANME abundance was positively correlated with these variables (Fig. 3E). Interestingly, across the coastal wetland types we investigated, salinity had little effect on methanogen functional type prevalence (Fig S7). *Methanosaeta* was the dominant genus consistent with previous reports in other freshwater wetland types and equally abundant in surface and deeper soils (Angle et al. 2017). Also, the family Methanomicrobiaceae has been detected in terrestrial wetlands and is often the dominant methane producer in Arctic wetlands (Wagner and Liebner 2010).

We expected a higher incidence of methylotrophs at higher salinities (Kallistova et al. 2020) partially due to increased availability of substrates such as trimethylamine (Bueno de Mesquita et al. 2023), which can be excreted by *Spartina* (Yuan et al. 2019). We note that the observed patterns are not necessarily due to the lower salinity range of our highest saltwater marsh, as other mesohaline wetlands have exhibited such patterns (Capooci et al. 2024). Our findings indicate that current theories on the distribution and potential importance of methanogen types across wetlands require further scrutiny and may not be universal across wetland salinity gradients. We suggest that other underappreciated factors like vegetation (Bansal et al. 2020; Bueno de Mesquita et al. 2024), salinity perturbation frequency (Kallistova et al. 2020; Hartman et al. 2024), eutrophication and resulting deoxygenation (Bonaglia et al. 2024; Żygadłowska et al. 2024), organic matter availability (Keneally et al. 2024) or other chemical factors (Zhang et al. 2023) that ultimately affect redox concentrations and geochemical cycling at the ecosite level (Axis 2 in Fig. 2C, Fig. 3G, I), may instead be playing a more prominent role in modulating biogeographic patterns of methanogens across coastal wetlands, particularly those with low salinity concentration ranges where salinity exerts limited selective pressure and other environmental drivers become more influential (Wallenius et al. 2021).

### 4.3 Methanogen richness and abundance predict *in situ* methane concentrations

Previous work has noted a positive correlation between methanogen abundance and methane flux or porewater concentrations via qPCR, mcrA gene sequencing, and 16S rRNA gene sequencing in rice paddies, freshwater wetlands, and lakes (Pierangeli et al. 2021). The use of similar methods to predict porewater methane concentrations is rare, and results vary widely, with methanogen abundance being either a strong (Perryman et al. 2022) or poor predictor (Rey-Sanchez et al. 2019). Though much more costly than 16S metabarcoding, *mcrA* expression has been shown to be a stronger predictor of *in situ* soil methane (Angle et al. 2017; Bechtold et al. 2025; Paul et al. 2026). This variability is expected, as many microbial taxa may be present but naturally dormant (Bickel and Or 2021) or inactive due to environmental stressors such as saltwater intrusion (Wen et al. 2019). Thus, while our results reveal methanogens as potential predictors of methane flux, given the variability in predictions across other sites using similar methods, further studies should include more activity-based measurements, such as metatranscriptomics, which directly account for methane production and oxidation and yield more consistent results.

### 4.5 Methanotroph relative abundance is highest in surface waters

We observed that methanotrophic bacteria were similarly abundant across sediment and water samples in the saltwater marsh. ANME1-b were abundant and discriminant of this site (Table S8), which is unsurprising as anaerobic oxidation of methane is a low-energy process with many members adapted to saline, high-sulfate wetlands where they may outcompete other methanotrophs (Kevorkian et al., 2021) while *Methylobacter* and *Methyloceanibacter* exhibited relatively high abundance across all saltwater marsh samples (Fig. 4). Notably, *Methylobacter* and other Type I methanotrophs have been shown to have the highest *pmoA* transcription and methane oxidization potential in freshwater lakes (Biderre-Petit et al. 2011), permafrost soils (Rainer et al. 2020), and other type of freshwater wetlands (Smith et al. 2018). While *Methylobacter* were only at high relative abundance in the saltwater marsh, this wetland had relatively high porewater methane concentrations and the lowest measured flux; thus, we hypothesize that *Methylobacter* could be the dominant contributor to the methane buffering capacity of this wetland.

The swamp soils, which had the highest methane porewater concentrations, were dominated by two Methylomirabilaceae genera, *Ca. Methylomirabilis* and Sh765B-TzT-35. Other *Methylomirabilis* species, including the denitrifier *M. oxyfera* (Yao et al. 2024), have been shown to be present and active in wetlands and freshwater lakes (Su et al. 2023). Though we did not measure denitrification rates or nitrous oxide fluxes for these soils, the presence of these taxa in the swamp suggests another possible source of greenhouse gas emissions from these wetlands. Together, these results suggest that the decoupling observed between porewater methane concentrations and methane fluxes may be explained by the methane filtering potential of methanotrophs in the water column (Fig. 6).

**Figure 6.**
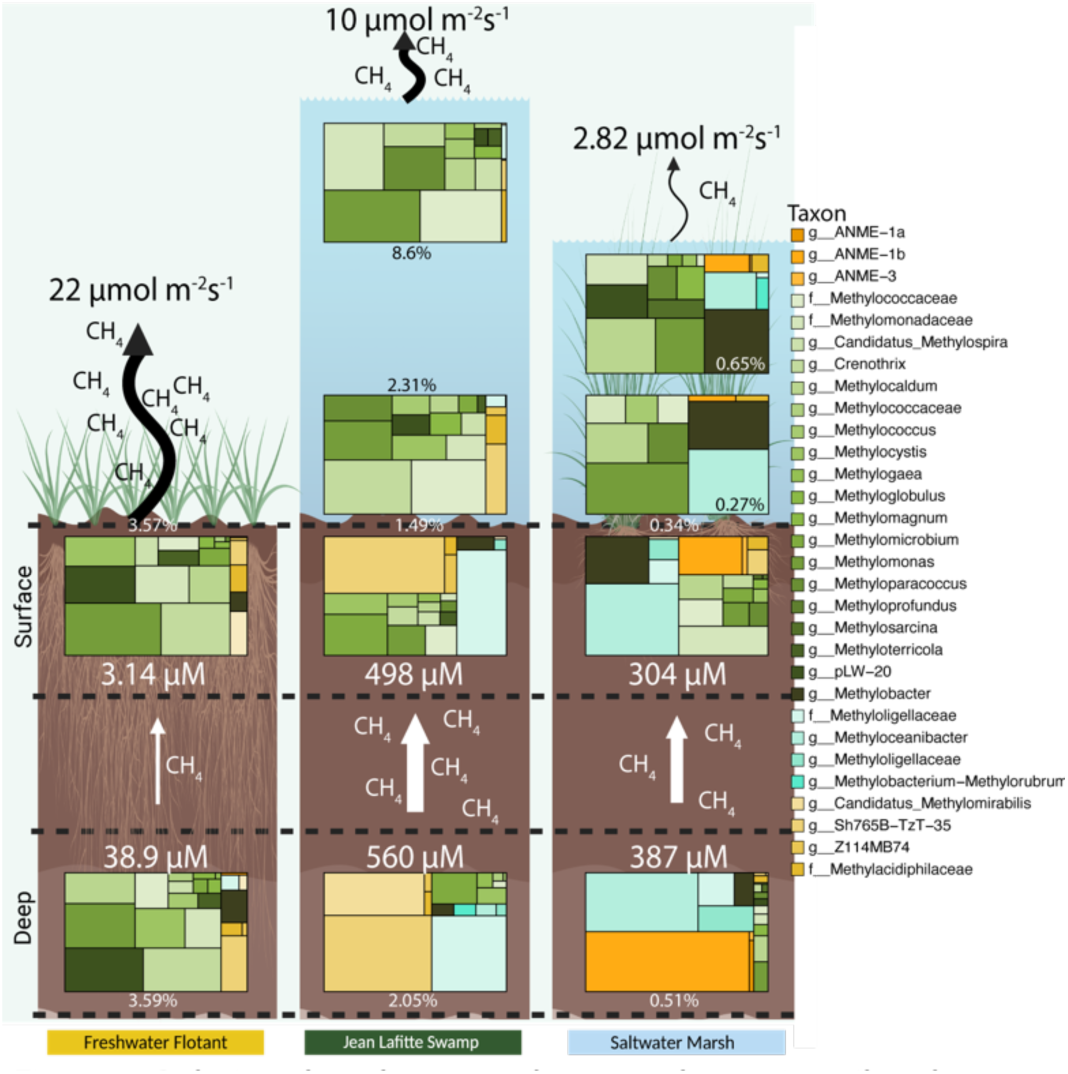
Relative abundance of methanotroph taxa in soil and water samples of each wetland. Percentages near each treemap denote the average relative abundance of methanotrophs among samples in each compartment. Colors within treemaps denote taxonomy, with shades of the same color representing methanotroph type. Black arrows and text above soil and water represent average methane flux per wetland. White arrows and text denote average methane porewater concentrations in surface (0-10cm) and deep (20-30cm) soils per wetland. Created in https://BioRender.com

## 5. Conclusions

We measured methane concentrations in the soil profile at different depths, calculated methane fluxes, and characterized the microbial community in soil and water samples in three coastal Louisiana wetlands along a salinity gradient. These included, by order of lowest to highest salinity, a freshwater flotant marsh, a low salinity swamp, and a mesohaline saltwater marsh. While accounting for ecosites with different microtopography and vegetation within each wetland, we observed significant methane concentrations and emissions variation among wetlands and ecosites. Fluxes were generally larger at the flotant marsh and decreased with salinity, while porewater concentrations were the lowest at the flotant marsh. Methanogen and methanotroph richness and abundance predicted methane concentrations across all wetland types. The core soil methane-cycling taxa were two methanogens: the obligate acetoclastic *Methanosaeta* and hydrogenotrophic Methanomicrobiaceae. Despite the lack of a single identifiable mechanism, methanogen and methanotroph community composition and functional type enrichment varied among wetlands, suggesting that the complex suite of environmental variables at each site select for different methane cycling taxa.

Understanding the distribution of and controls on microbial methane production from wetlands is critical to global methane emission forecasting, particularly in response to changing environmental conditions such as rising temperatures and saltwater intrusion. Our observations show that simplistic common perceptions about the biogeography of methanogens along salinity gradients (e.g., enriched methylotrophs, negatively impacted hydrogenotrophs) warrant further functional investigation across coastal wetland systems. These results also highlight the need to examine microbial communities across the terrestrial-aquatic interface and not consider any one compartment in isolation. The biological framework presented here can be used to generate knowledge for increased realism in predictive, process-oriented models of methane fluxes.

## Acknowledgments

We thank Tyson Claffey and Richard Wolfe for Colorado State University server management and Rebecca Daly for Colorado State University laboratory management. Grammarly was used to correct grammatical errors and improve the clarity of text. This work was partially supported by awards from the U.S. Department of Energy (DOE) Office of Science, Office of Biological and Environmental Research (BER) grants DE-SC0023084 (DKMF, SKJ, JBE, BBM, GB, EW, MAB, KCW, JAV), DE-SC0021067 (GB, KCW, JAV), and DE-SC0022972 (JAV). Amplicon sequencing was performed at the Microbial Community Sequencing Lab, University of Colorado Boulder.

